# Small Molecule Screen Identifies Non-Catalytic USP3 Chemical Handle

**DOI:** 10.1101/2023.03.01.530657

**Authors:** Mandeep K. Mann, Esther Wolf, Madhushika Silva, Haejin Kwak, Brian Wilson, Derek J. Wilson, Rachel J. Harding, Matthieu Schapira

## Abstract

Zinc-finger ubiquitin binding domains (ZnF-UBDs) are non-catalytic domains mostly found in deubiquitylases (DUBs). They represent an underexplored opportunity for the development of deubiquitylase-targeting chimeras (DUBTACs) to pharmacologically induce the deubiquitination of target proteins. We have previously shown that ZnF-UBDs are ligandable domains. Here, a focused small molecule library screen against a panel of eleven ZnF-UBDs led to the identification of **59**, a ligand engaging the ZnF-UBD of USP3 with a K_D_ of 14 µM. The compound binds the expected C-terminal ubiquitin binding pocket of USP3 as shown by hydrogen-deuterium exchange mass spectrometry experiments and does not inhibit the cleavage of K48-linked di-ubiquitin by USP3. As such this compound could serve as a chemical starting point to develop bifunctional DUBTACs recruiting USP3 for targeted deubiquitination.

**Table of contents graphic:** 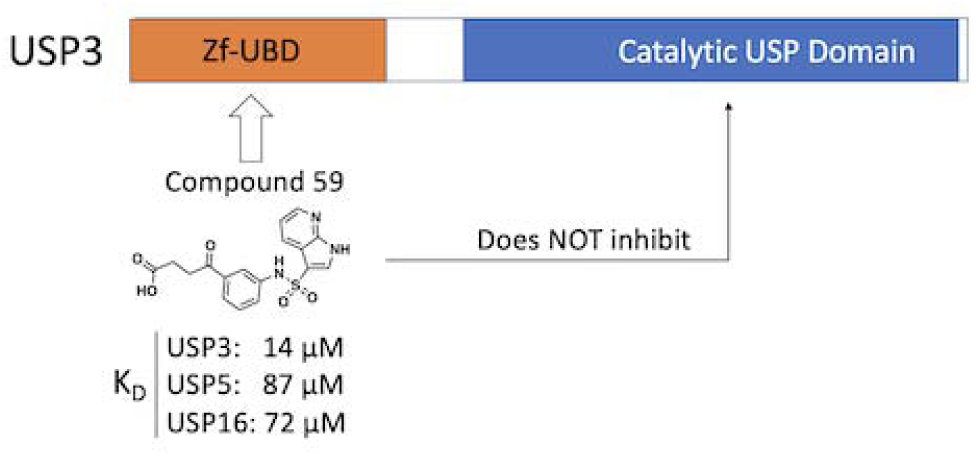

## INTRODUCTION

Proximity-induced pharmacology is an expanding field in drug discovery, in which post-translational modifications can be induced through the recruitment of protein-modifying enzymes to target proteins. Mechanistic proof-of-concept was established with proteolysis targeting chimeras (PROTACs) that induce ubiquitination and proteasomal degradation of a target protein via the recruitment of an E3 ubiquitin ligase. Conversely, the ability to pharmacologically rescue proteins from proteasomal degradation can allow the investigation of protein function and may have therapeutic benefits in disease settings where a protein is aberrantly degraded. Bifunctional molecules presenting a chemical handle binding a non-catalytic pocket of a deubiquitylase (DUB), an enzyme that removes ubiquitin groups, may enable targeted protein deubiquitination and subsequent target stabilization. The first reported deubiquitylase-targeting chimera (DUBTAC) recruited OTUB1, a K48 ubiquitin specific DUB^1^. The compound was composed of a covalent ligand binding an allosteric site of OTUB1, linked to a drug that binds a mutant cystic fibrosis transmembrane conductance regulator (CFTR), resulting in stabilized CFTR protein levels in human cystic fibrosis bronchial epithelial cells and restored chloride channel function^1^.

The zinc-finger ubiquitin binding domain (ZnF-UBD), is a non-catalytic structural module found in twelve USPs (USP3, USP5, USP13, USP16, USP20, USP22, USP33, USP39, USP44, USP45, USP49, USP51)^2–4^ with a unique fold that primarily binds the C-terminal RLRGG motif of ubiquitin (Ub). The ZnF-UBD is also found in the lysine deacetylase HDAC6 and BRAP, a BRCA-1 associated protein and Ub ligase^5–8^ (**Figure 1A**). While ZnF-UBDs generally engage Ub with a low micromolar binding affinity, this function is lost in some USPs (USP13, USP20, USP22, USP33, USP39 and USP51) due to substitutions of key ubiquitin-coordinating residues^3, 9–11^ (**Figure 1B**). We previously demonstrated the ligandability of the ZnF-UBD with low micromolar ligands for the ZnF-UBD of USP5 and a <100 nanomolar ligand for HDAC6^12–15^ (**Figure 1AC**). The function of ZnF-UBDs and their role in catalytic activity of DUBs is poorly understood. We have shown that ligands targeting the ZnF-UBD of USP5 inhibit its catalytic activity^12, 13^ and therefore cannot be used as chemical handles for the development of USP5-recruiting DUBTACs, but the same may not be true for small molecules targeting the ZnF-UBD of other USPs.

**Figure 1.**
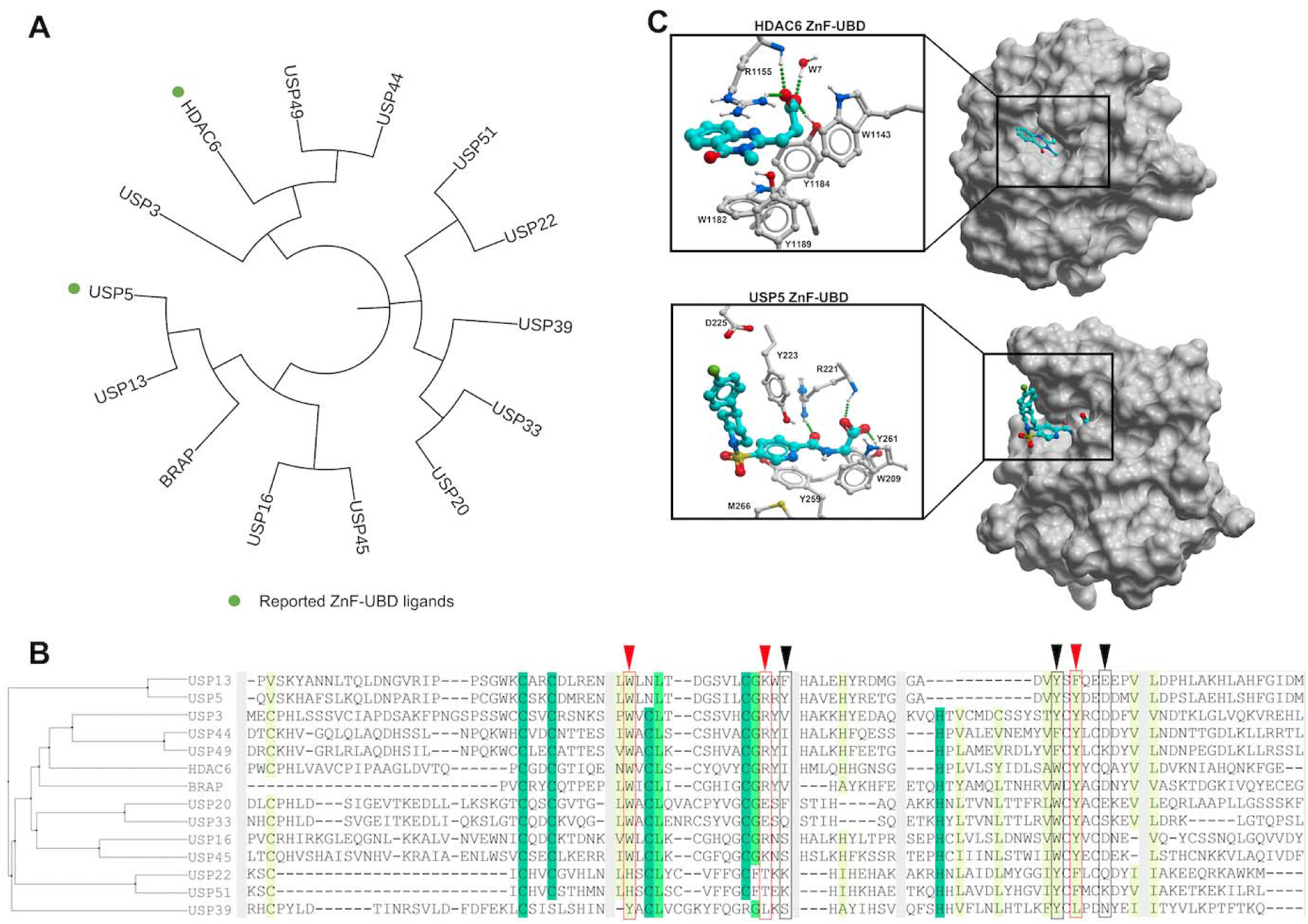
Human ZnF-UBDs: A) (Left) Dendrogram of full-length human proteins that contain a ZnF-UBD. The dendrogram was created by first doing a multiple sequence alignment using Clustal Omega^16^, followed by tree generation with iTol^17^. Reported ZnF-UBD ligands are highlighted with green circles B) Sequence alignment of human ZnF-UBDs. Conserved residues are indicated with shading. USP5 residues that interact with ubiquitin are indicated with arrowheads and boxes (red: expected to be essential for ubiquitin binding; black: not essential). C) Co-crystal structures of HDAC6 and USP5 ZnF-UBD in complex with low micromolar ligands (PDB: 6CED, 7MS7). Hydrogen bonds are displayed as dashed lines.

Here, we report the discovery and characterization of USP3 ZnF-UBD ligands. Screening of a small, focused chemical library against a panel of ZnF-UBDs led to the discovery of a selective USP3 ligand that does not inhibit the catalytic function of USP3. We also present a preliminary exploration of the structure activity relationship (SAR) of the chemical series. Our work provides a framework for the development of more potent USP3 ZnF-UBD ligands as chemical handles for DUBTACs, and as tools to study the function of the USP3 ZnF-UBD in health and disease.

## MATERIALS AND METHODS

### Cloning, Protein Expression and Purification

cDNAs encoding Ub^1–76^, Ub^1–73^, USP5^171–290^, and HDAC6^1109–1215^ were cloned as previously described^12–15^. DNA encoding USP3^1–131^, USP16^25–185^, USP20^1–141^, USP33^29–134^, USP39^84–194^, USP49^1–115^, USP51^176–305^ and BRAP^304–390^ was sub-cloned into a modified vector encoding an N-terminal AviTag for biotinylation and a C-terminal His6 tag (p28BIOH-LIC), while USP13^183–307^ was sub-cloned into a vector with an N-terminal His-tag and TEV protease cleavage site, and a C-terminal biotinylation sequence (pNICBIO2). USP3^1–520^ was sub-cloned into a pFBOH-LIC vector and overexpressed in sf9 cells, where cultures were grown in HyQ SFX Insect Serum Free Medium (Fisher Scientific) to a density of 4 × 10^6^ cells/mL and infected with 10 mL of P3 viral stock media per 1 L of cell culture. Cell culture medium was collected after 4 days of incubation in a shaker at 27LC. Proteins were purified as before^12–15;^ briefly, all proteins were purified by metal affinity chromatography, gel filtration and ion exchange chromatography. The final concentration of purified proteins was measured by UV absorbance at 280 nm. Protein identity was confirmed by mass spectrometry and purity was assessed by SDS-PAGE.

### Surface Plasmon Resonance Assay

Studies were performed with a Biacore^TM^ T200 (GE Health Sciences) at 20LC. Approximately 3000-6000 response units (RU) of biotinylated ZnF-UBDs were captured to flow cells of a streptavidin-conjugated SA chip per manufacturer’s protocol, and an empty flow cell used for reference subtraction. Serial dilutions were prepared in 20 mM HEPES pH 7.4, 150 mM NaCl, 1 mM TCEP, 0.005% Tween-20 (v/v), 1% DMSO (v/v). K_D_ determination experiments were performed using multi-cycle kinetics with 60 second contact time, and 60 second dissociation time at a flow rate of 30 µL/min at 20 LC. K_D_ values were calculated using steady state affinity fitting with the Biacore^TM^ T200 evaluation software (GE Health Sciences).

### Ubiquitin Rhodamine110 Assay

Experiments were performed in a total volume of 60 µL in 384-well black polypropylene microplates (Grenier). Fluorescence was measured using a Biotek Synergy H1 microplate reader (Biotek) at excitation and emission wavelengths of 485 nm and 528 nm, respectively. Ligands were prepared in 20 mM Tris pH 7.5, 125 mM NaCl, 1 mM DTT, 0.01% TX-100 (v/v), 1% DMSO (v/v) for two-fold dilution titration series. 500 nM USP3^1–520^ and 200 nM ubiquitin-rhodamine110 (UBPBio) were added to each well. Following a 1 minute centrifugation at 250 g, fluorescence readings were immediately taken for ten minutes. The data was analyzed with GraphPad Prism 8.2.0.

### Ub2K48 Cleavage Assay

70 pmoles each of USP3 full-length (USP3^1–520^) or USP3 ZnF-UBD (USP3^1–131^) were incubated in 10 mM HEPES pH7.5, 150 mM NaCl, 3 mM EDTA, 0.05% Tween-20, 1 mM TCEP with or without compound 59 at 10-fold molar excess, for 1 hour at room temperature. 60 pmoles of Ub2K48 (Boston Biochem) was added to all reactions, bringing the final volume to 10 μL and reactions were incubated for 1 hour at room temperature. Reactions were stopped by addition of SDS-PAGE loading dye to a final concentration of 1X and incubation at 95°C for 5 minutes. Reactions were analysed by Coomassie stained SDS-PAGE using NuPAGE 4-12% Bis-Tris gel Invitrogen).

### Hydrogen-Deuterium Exchange Mass Spectrometry (HDX-MS)

To begin, three samples were prepared and incubated for 30 min at 0 °C: 7.5 μM USP3 ZnF-UBD (apo), 7.5 μM USP3 ZnF-UBD and 75 μM **59** (1:10 complex), 7.5 μM USP3 ZnF-UBD and 150 μM **59** (1:20 complex). All samples were then held at 0 °C prior to mixing 8 μL of sample with 52 μL of buffered D_2_O (10 mM Phosphate Buffer pD 7.5, 150 mM NaCl) to yield a final D_2_O concentration of 87%. The HDX reaction was allowed to take place for 15, 60, or 600 seconds at 20°C. Then, 50 μL of the HDX reaction was quenched for 1 min at 0 °C with 50 μL Quench buffer (100 mM Phosphate Buffer, pH 2.5) to stop the HDX reaction. Next, 50 μL (25 pmol) of the quenched sample was loaded onto a mixed Nepenthensin-2 Pepsin (1:1) column (Affipro) and desalted (ACQUITY UPLC BEH C18 VanGuard Pre-column, 130Å, 1.7 µm, 2.1 mm × 5 mm, Waters) using Mobile Phase A (0.1% formic acid in water) at 200 μL/min for 3 mins. The peptides were then reverse-phase separated (ACQUITY UPLC BEH C18 Column, 130Å, 1.7 µm, 2.1 mm × 50 mm, Waters) at 40 μL/min using a gradient from 5 to 35 % Mobile Phase B (0.1% formic acid in acetonitrile). Eluted peptides were electrosprayed into the Select Series Cyclic IMS (Q-IMS-TOF, Waters) while Leu-Enk was used for lockspray. Fragmentation was conducted using collision-induced dissociation (CID) in HDMS^e^ mode with a collisional energy ramp from 20 to 29 V (in the transfer cell). To obtain undeuterated peptides, the above steps were followed except in place of deuteration buffer, an equilibration buffer was used (10 mM Phosphate Buffer pH 7.5, 150 mM NaCl). HDX was automated using the ACQUITY UPLC M-Class System with HDX Technology (Waters) and PAL3 liquid handling (CTC Analytics AG). Peptide identification was conducted using ProteinLynx Global Server 3.0.3 (PLGS, Waters). HDX analysis and visualization was conducted using DynamX 3.0 (Waters) and PyMOL 2.5.0.

### Molecular Modeling

The model of USP3 predicted by AlphaFold was downloaded from https://alphafold.ebi.ac.uk/ and superimposed to the crystal structure of USP5 ZnF-UBD in complex with a chemical analog of **59**. Compound **59** was then docked to the corresponding pocket using a grid representation of the receptor with ICM (Molsoft, San Diego). The system was further relaxed with 100 ns molecular dynamics (MD) simulations with ICM using OpenMM ^18^. Six independent simulations did not all converge to the same pose, reflecting the challenge of docking to AlphaFold structures. The binding pose shown in **Figure 4** was from one of the most stable simulations (**Figure S2**) and is provided as an example. All binding poses where at the same site and occluded residues 91-95.

### Hit Expansion

Substructure search was run against the Enamine REAL database (Enamine Ltd., 2022). Compounds were clustered in ICM (Molsoft) ^19^ and selected to be purchased based on availability, cost, and chemical substitutions. The purity of all compounds in **Table 1** was confirmed to be at least 95% as measured by LCMS.

**Table 1.**
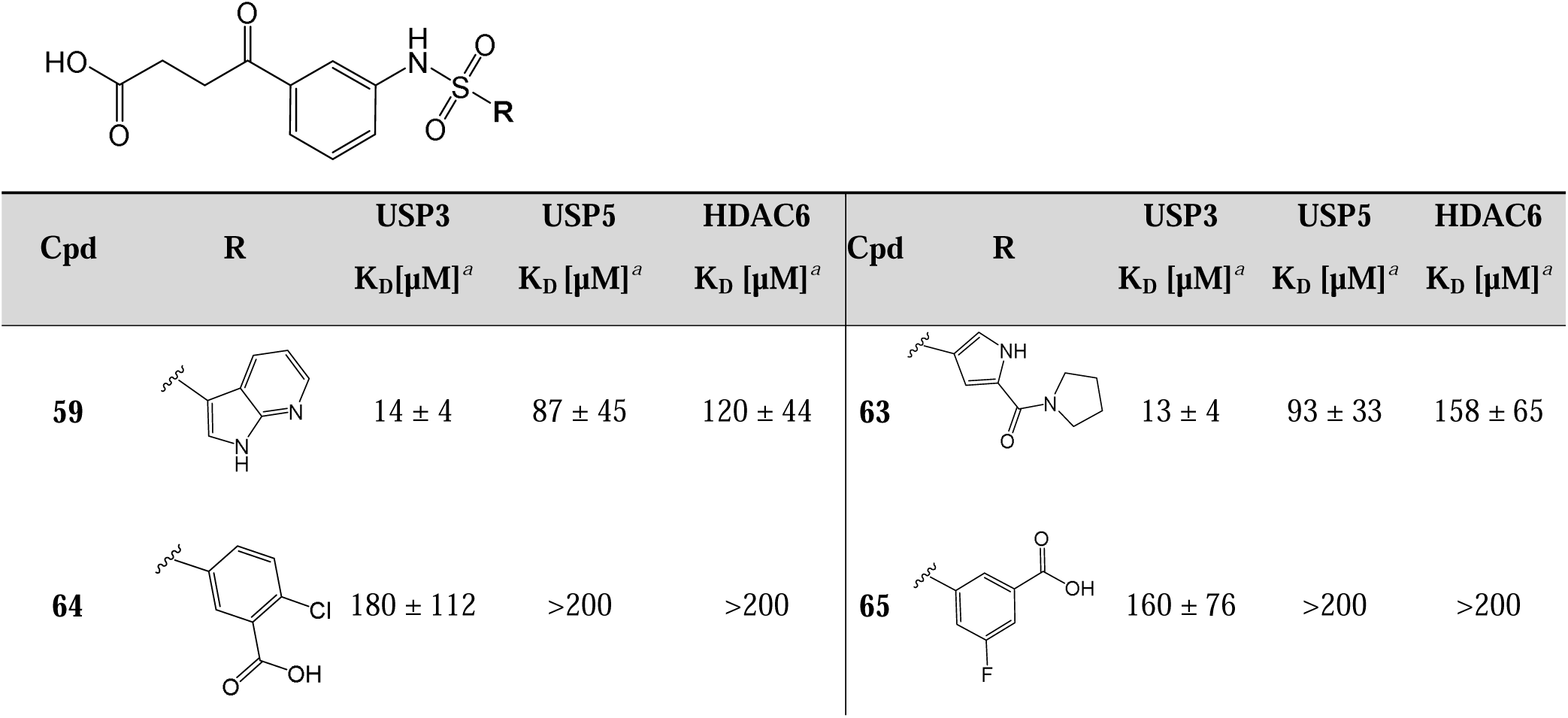
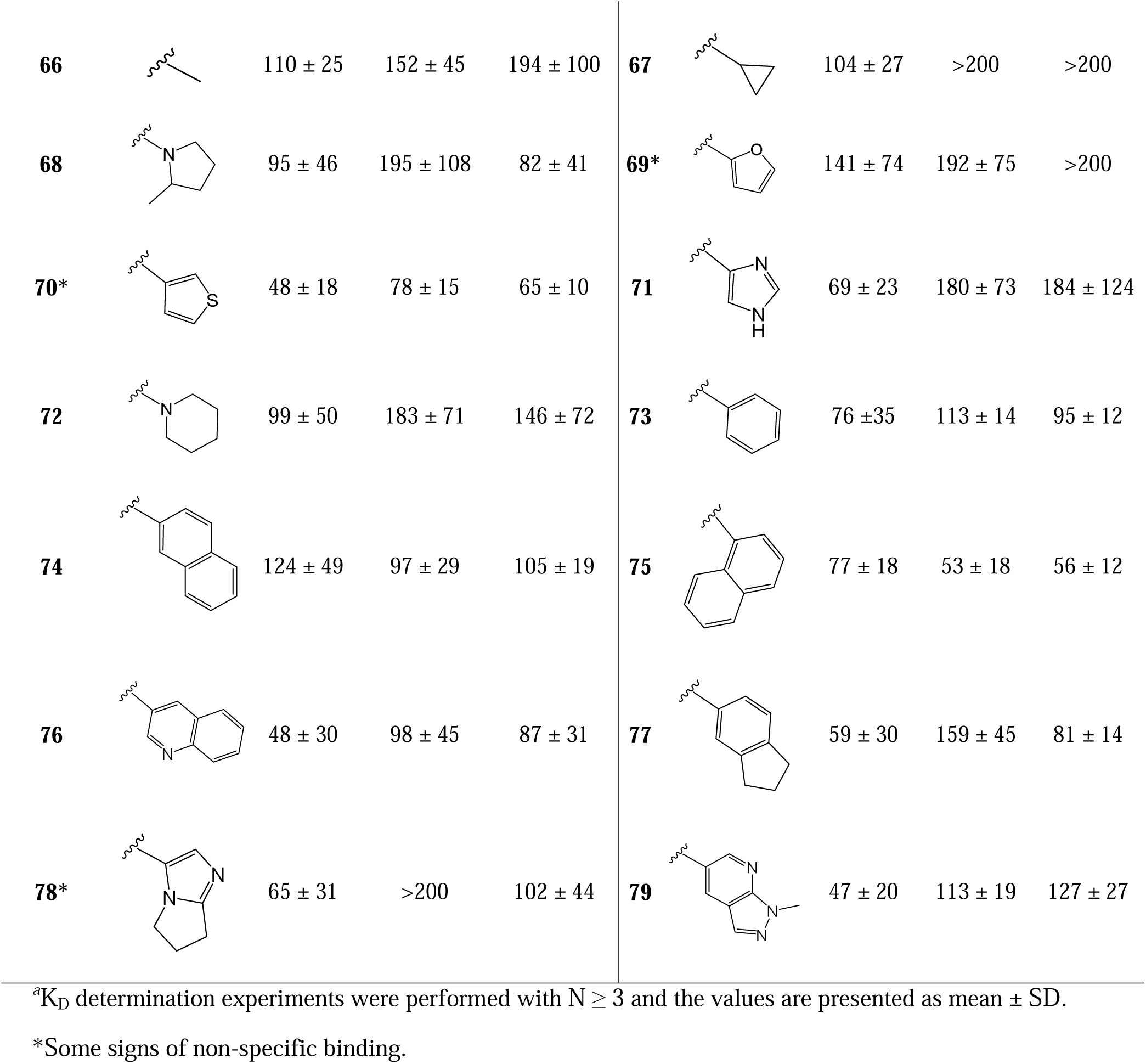
In-house and Enamine REAL analogs of **59**

|  |  |  |  |  |  |  |  |  |  |
| --- | --- | --- | --- | --- | --- | --- | --- | --- | --- |
| Cpd | R | USP3<br>$K_D[\mu\text{M}]^a$ | USP5<br>$K_D[\mu\text{M}]^a$ | HDAC6<br>$K_D[\mu\text{M}]^a$ | Cpd | R | USP3<br>$K_D[\mu\text{M}]^a$ | USP5<br>$K_D[\mu\text{M}]^a$ | HDAC6<br>$K_D[\mu\text{M}]^a$ |
| **59** | | $14 \pm 4$ | $87 \pm 45$ | $120 \pm 44$ | **63** | | $13 \pm 4$ | $93 \pm 33$ | $158 \pm 65$ |
| **64** | | $180 \pm 112$ | >200 | >200 | **65** | | $160 \pm 76$ | >200 | >200 |
| **66** |  | 110 ± 25 | 152 ± 45 | 194 ± 100 | **67** |  | 104 ± 27 | >200 | >200 |
| **68** |  | 95 ± 46 | 195 ± 108 | 82 ± 41 | **69\*** |  | 141 ± 74 | 192 ± 75 | >200 |
| **70\*** |  | 48 ± 18 | 78 ± 15 | 65 ± 10 | **71** |  | 69 ± 23 | 180 ± 73 | 184 ± 124 |
| **72** |  | 99 ± 50 | 183 ± 71 | 146 ± 72 | **73** |  | 76 ± 35 | 113 ± 14 | 95 ± 12 |
| **74** |  | 124 ± 49 | 97 ± 29 | 105 ± 19 | **75** |  | 77 ± 18 | 53 ± 18 | 56 ± 12 |
| **76** |  | 48 ± 30 | 98 ± 45 | 87 ± 31 | **77** |  | 59 ± 30 | 159 ± 45 | 81 ± 14 |
| **78\*** |  | 65 ± 31 | >200 | 102 ± 44 | **79** |  | 47 ± 20 | 113 ± 19 | 127 ± 27 |

<sup>a</sup>K<sub>D</sub> determination experiments were performed with N ≥ 3 and the values are presented as mean ± SD.
\*Some signs of non-specific binding.

## RESULTS

### Discovery of a ligand binding the ZnF-UBD of USP3

Following our observation that an aliphatic chain ending with a carboxylic acid mimicking the endogenous substrate of the ZnF-UBD of HDAC6 and USP5 was essential for binding, we compiled a focused library of 670 molecules bearing this moiety from our in-house collection to target the Ub-binding ZnF-UBDs of USP3, USP5, USP13, USP16, USP20, USP22, USP33, USP39, USP49, USP51, BRAP and HDAC6. Compounds were grouped into 62 clusters with ICM (Molsoft) based on their chemical similarity, and the binding of 62 representative molecules to our panel of 11 ZnF-UBDs was measured in a surface plasmon resonance (SPR) assay. Ub and a C-terminal RGG-deleted Ub construct were used as positive and negative controls, respectively (**Figure 2A, Table S1 & S2)**. As expected, no binding to Ub was observed for USP13, USP20, USP33, USP39 and USP51, but low micromolar binding was observed for USP3, USP5, USP16, BRAP and HDAC6. No binding was observed to RGG-deleted Ub by any ZnF-UBD protein. Interestingly, no Ub binding was observed for USP49, despite conservation of Ub-binding residues^20, 21^.

**Figure 2.**
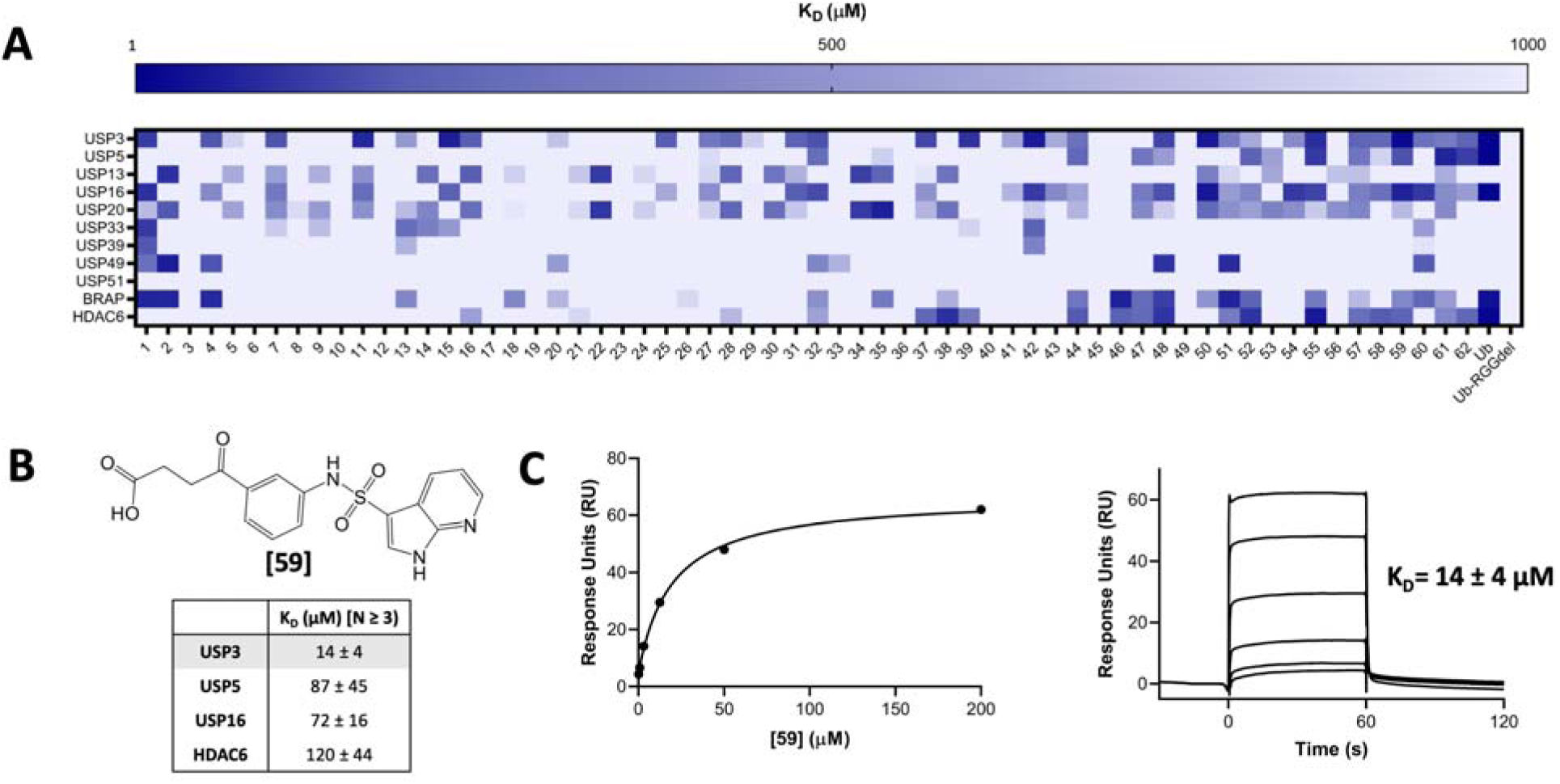
A small molecule screen identifies ZnF-UBD hits. A) A heat map showing binding of 64 ligands to 11 ZnF-UBDs by SPR. A 4-fold 6-point dilution series beginning at 200 µM wa used for K_D_ determination with N=1. B) Summary of binding data for compound **59** by SPR, N≥3. C) Representative SPR binding curve from steady-state fit analysis and sensorgrams for USP3 ZnF-UBD and **59**. A K_D_ of 14 ± 4 was obtained from the average of seven independent measurements.

Many ligands from the screen bound USP3 but few showed selectivity for USP3 over other ZnF-UBDs. Compound **59** was identified and confirmed as a promising USP3 hit and was more than 5-times selective for USP3 over USP5, USP16, and HDAC6 ZnF-UBDs (K_D_: USP3 = 14 ± 4 µM; USP5 = 87 ± 45 µM; USP16 = 72 ± 16 µM; HDAC6 = 120 ± 44 µM) (**Figure 2BC**, **Table 1**).

### Compound 59 does not perturb the enzymatic activity of USP3

We next used a fluorogenic ubiquitin rhodamine assay to test whether **59** inhibited USP3 DUB catalytic activity. We found that the enzyme remains fully active even in the presence of 1 mM compound **(Figure 3A**). This is in contrast with USP5, where ligands targeting the ZnF-UBD inhibit catalytic activity^13^. To verify that binding of C-terminal ubiquitin tail to the ZnF-UBD of USP3 was not necessary for the cleavage of endogenous substrates, we used a gel-based assay to monitor the cleavage of a K48-linked diubiquitin substrate in the absence and presence of **59**. Again, we saw no sign of inhibition of deubiquitinase activity, suggesting that USP3 ZnF-UBD targeting ligands could be used as chemical starting points for USP3-recruiting DUBTACs (**Figure 3B**).

**Figure 3:**
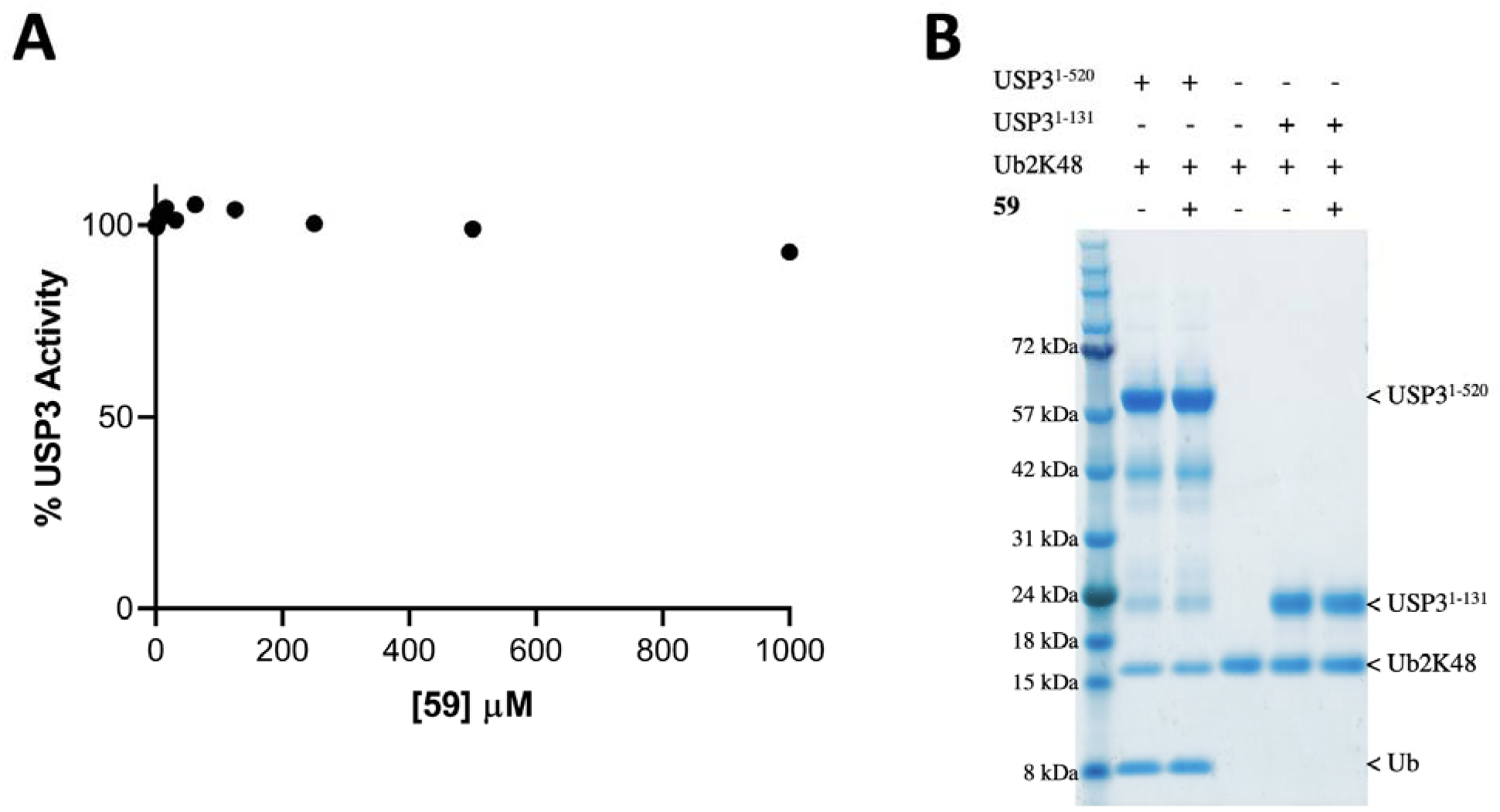
59 does not inhibit USP3 catalytic activity. A) UbRho110 assay (N=2). Fluorescence signal is normalized to control (no compound, DMSO only). B) Cleavage of Ub2K48 in the absence or presence of **59 (**70 µM final concentration).

**Figure 4.**
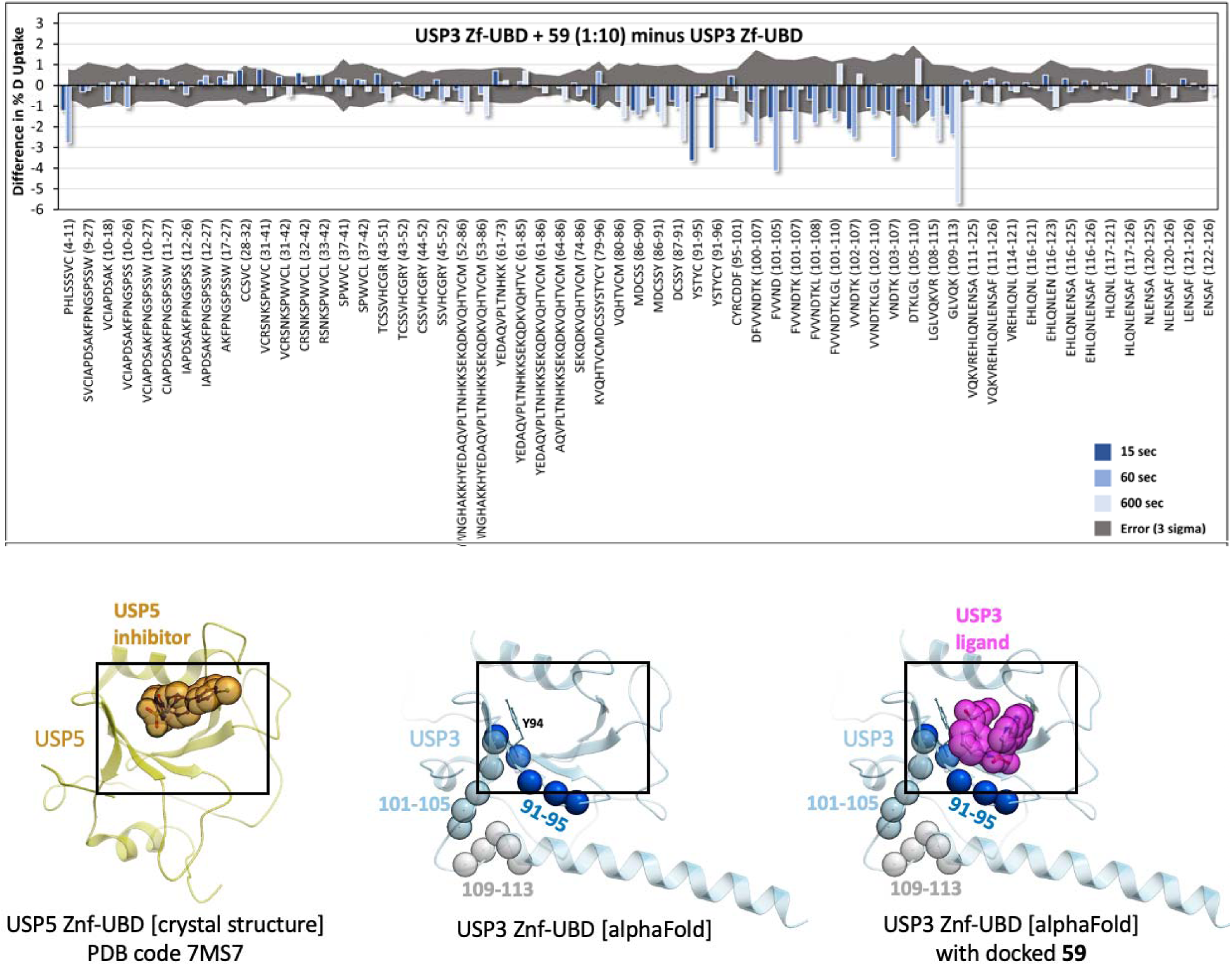
Mapping of the ligand binding site by HDX-MS. Top: the difference in deuterium fractional uptake (%) between USP3 ZnF-UBD + **59** (1:10) and USP3 ZnF-UBD is plotted as a function of each timepoint per peptide. The timepoints are 15 s (yellow), 60 s (orange), and 600 s (red). The error (3 times the propagated error) is shown as grey area. If the bars exceed this grey area, they are statistically significant. Bars pointing down indicate a decrease in deuterium uptake in the presence of the ligand. Bottom: HDX signals observed at 15 s (dark blue), 60 s (light blue) and 600 s (grey) are highlighted are spheres on an AlphaFold prediction of the USP3 ZnF-UBD structure (center). A crystal structure of the homologous USP5 ZnF-UBD bound to a ligand (PDB ID: 7MS7) (left) and a model of **59** docked to the USP3 ZnF-UBD AlphaFold model prediction (right) are shown as references.

### Compound 59 binds the C-terminal ubiquitin binding site of USP3

The structure of USP3 ZnF-UBD is not available from the PDB and despite extensive crystallization screening efforts, we were unable to solve it by X-ray crystallography, in either its apo form or in complex with ubiquitin or **59**. To identify the binding site of **59** and characterize the binding event’s corresponding allosteric effects, hydrogen-deuterium exchange mass spectrometry (HDX-MS) was used. A sequence coverage of over 90% was obtained for USP3 ZnF-UBD with a peptide per amino acid redundancy of 5.24 (see supplementary **Table S3** and **Figure S1**).

HDX-MS analysis was performed for USP3 ZnF-UBD alone and in complex with **59** (1:10 and 1:20) for 15 s, 60 s, and 600 s at 20 °C (**Figures 4** and **S1, S2**). Given the 14 μM dissociation coefficient (K_D_), the bound fractions in the 1:10 and 1:20 states upon D_2_O buffer addition were expected to be 40.1% and 58.1%, respectively. The HDX data described here are differential (deuterium uptake of bound state minus unbound state), in technical triplicate (n=3), and both the 1:10 and 1:20 ligand ratios show consistent results (effectively n=6). To be considered statistically significant, differences must have exceeded triple the propagated error. Complexation resulted in significant attenuation of deuterium uptake rates in 20 out of 59 peptides, all localized to the same region (**Figure 4**). At the earliest timepoint (15 s), distinct high-intensity decreases were observed at residues 91-95 (YSTYC) which quickly disappear and are absent at the 1 and 10 min timepoint. The next set of high intensity decreases appear at 1 min for residues 5-10 (HLSSSVC) and 102-107 (VVNDTK). Finally, residues 110-113 (LVQK) exhibit slowly evolving decrease in deuterium uptake, whose magnitude becomes the dominant difference signal at the longest measured timepoint (10 mins).

Together, these data support a direct binding event at residues 91-95 (as discussed below), which superimpose with the binding pocket of an analogous compound co-crystallized in complex with the ZnF-UBD of USP5 (**Figure 4**) ^13^. A predicted model of **59** bound to this site shows that the compound is expected to occlude residues 91-95 from solvent, in agreement with the decrease in deuterium exchange observed immediately at this site upon treatment with **59** (**Figure 4**).

### Some chemical analogs also bind USP3 ZnF-UBD

To validate the chemical template of **59**, three in-house analogs were tested for binding to the ZnF-UBD of USP3, USP5 and HDAC6, confirming that potency and selectivity varied with substituents attached to the sulfonamide group (**Table 1**, **63-65**). A substructure search of the Enamine REAL database for analogs of **59** and **63** (K_D_ = 13 ± 5 µM) resulted in 797 compounds, which were clustered with ICM (Molsoft). Fourteen diverse molecules were ordered and tested for binding to define a preliminary structure activity relationship (SAR) (**Table 1**, **66-79**). While more potent molecules were not identified, this limited SAR exploration showed that hetero-bicyclic rings with a nitrogen in 2-position of the sulfonamide linker are preferred (**59**, **63**, **76, 79**).

## DISCUSSION

USP3 is a nuclear DUB that primarily deubiquitinates and stabilizes targets for maintenance of genome stability, regulation of cell proliferation, DNA damage response, and the innate immune response^22–27^. There are currently no selective USP3 inhibitors, due to the high conservation of the catalytic USP domain. The ZnF-UBD of USP3 is required for its interaction with and deubiquitination of targets such as histone H2A and RIG-I^22, 25^.

Here, we identified a ligand of the USP3 ZnF-UBD that does not inhibit the enzymatic activity of full-length USP3. We were unable to solve the crystal structure of the USP3 in complex with **59** but used HDX-MS to map its binding site. While our HDX data indicate a direct binding event at a site known in the context of USP5 to recognize the C-terminal extremity of ubiquitin ^2^, and to be exploited by chemical analogs of **59** ^13^, we are also observing deuterium exchange at an adjacent site (**Figure 4**). Attenuated deuterium uptake kinetics in the presence of a ligand can arise from ‘direct’ effects (*i.e.*, new intermolecular hydrogen bonding or solvent exclusion resulting from direct contact between the protein and the ligand) or ‘induced’ effects (*i.e*., increased intramolecular hydrogen-bond stability upon binding, resulting in reduced sampling of a broader conformational ensemble accessible to the native, unbound protein) ^28–30^. HDX uptake kinetics can therefore provide insight on whether an observed decrease in deuterium uptake in a particular region is a direct or allosteric effect of ligand binding ^31^. Briefly, this is because direct effects are a function of the rate of binding equilibration (i.e., k_on_ + k_off_), which in this case will cause an ‘early’ time of maximum difference in the HDX data that rapidly diminishes due to the high k_off_ of **59** (as observed in the SPR sensorgrams, see **Figure 2C**). Indeed, the HDX data indicates that **59** makes direct contact at residues 91-95, which uniquely exhibits the expected direct effect difference kinetics profile. Notably, this agrees exactly with a critical π stacking in the docked model between the acetophenone group of **59** and Y94 (**Figure 4**), corresponding to π-stacking of the crystallized USP5 ligand with Y259. All other significant decreases in uptake, corresponding to residues 102-107, and 110-113 are near the predicted binding pocket, but exhibit slower uptake kinetics, indicative of allosteric induced effects. All HDX differences observed localize to one region of USP3, highlighting the same pocket verified in an analogous structure, PDB 7MS7. Here, the USP5 ZnF-UBD is bound to compound N-{5-[4-(4-chlorophenyl)piperidine-1-sulfonyl]pyridine-2-carbonyl}glycine (ZQ1), which shares a similar chemical scaffold to **59** ^13^. In the USP5 structure, there is a close π-stacking interaction between the pyridine moiety of ZQ1 and Y259, which is analogous to the HDX-supported **59**/USP3 Y94 interaction discussed above.

Compound **59** binds USP3 with a K_D_ around 15 µM, which would not be sufficient for a traditional occupancy-based competitor. Proximity inducing pharmacological agents such as PROTACs bind at the interface of proteins and exploit, rather than compete against protein-protein interactions. As such, they can sometimes be derived from chemical handles with relatively low affinity. As an example Apcin, a ligand binding the E3 ligase CDC20 with a K_D_ of 10 µM, was successfully used as a chemical handle for the development of a CDC20-recruiting PROTAC^32^.

In conclusion, compounds presented here may be good chemical starting points for the development of protein-protein interaction inhibitors that would antagonize binding of USP3 to some of its endogenous targets. Conversely, **59** or more potent compounds may serve as chemical handles for the development of bifunctional molecules that recruit USP3 to neo-substrates and induce their deubiquitination. In addition to the identification of **59**, the focused chemical library screen also identified hits against other ZnF-UBDs, such as USP16, USP49, and BRAP (**50**, **2**, **46**, respectively) which require further validation, but may be promising starting points for hit expansion.

## ASSOCIATED CONTENT

### Supporting Information

- Table S1: Compounds smiles strings and catalog numbers
- Table S2: Matrix Screen results
- Table S3: PLGS filtering parameters
- Figure S1: HDX sequence coverage of USP3^1–131^
- Figure S2: Differential HDX heatmaps
- Figure S3: Ligand RMSD over 100 ns MD simulation

## AUTHOR INFORMATION

### Author Contributions

The manuscript was written by M.K.M., E.W., D.J.W and M.S. All authors have given approval to the final version of the manuscript. M.K.M expressed and purified proteins, designed and optimized biophysical assays and tested compounds., B.W conducted analytical chemistry. E.W conducted and D.J.W supervised HDX experiments. H.K conducted molecular modeling simulations. R.J.H. and M.S. advised throughout the project.

### Notes

The authors declare no competing financial interest.

## ACKNOWLEDGMENTS

The Structural Genomics Consortium is a registered charity (no: 1097737) that receives funds from Bayer AG, Boehringer Ingelheim, Bristol Myers Squibb, Genentech, Genome Canada through Ontario Genomics Institute [OGI-196], EU/EFPIA/OICR/McGill/KTH/Diamond Innovative Medicines Initiative 2 Joint Undertaking [EUbOPEN grant 875510], Janssen, Merck KGaA (aka EMD in Canada and US), Pfizer and Takeda. We thank Dr. Vijayaratnam Santhakumar for his advice and Albina Bolotokova for compound management.

## ABBREVIATIONS

BRAP, BRCA-1 associated protein; DMSO, dimethylsulfoxide; DUB, deubiquitylase; DUBTAC, deubiquitylase targeting chimera; HDAC6, histone deacetylase 6; HDX: hydrogen deuterium exchange; K_D_, dissociation constant; K48, lysine 48; MD, molecular dynamics; PROTAC, proteolysis targeting chimera; SAR, structure activity relationship; SD, standard deviation; SPR, surface plasmon resonance; Ub, ubiquitin; USP, ubiquitin specific protease; ZnF-UBD, ubiquitin-specific processing protease zinc-finger

