## Supplementary information for "Small Molecule Screen Identifies Non-Catalytic USP3 Chemical Handle"

- Table S1: Compounds smiles strings and catalog numbers …………. p.2
- Figure S3: Ligand RMSD over 100 ns MD simulation ……………....p.7

**Table S1: Compounds smiles strings and catalog numbers**

| Compound # | Catalog_Number | SMILES |
| --- | --- | --- |
| 1 | STL102634 | C(C([O-])=O)c1cccnc1 |
| 2 | STK256632 | C(Cn1cccn1)C([O-])=O |
| 3 | STK661679 | C(Cc1ccno1)C([O-])=O |
| 4 | Z57127332 | C(Cc1ccccc1)C(O)=O |
| 5 | STK802016 | C(C([O-])=O)Nc1ncccn1 |
| 6 | IF_02_176 | C(Cc1cnccn1)C(O)=O |
| 7 | SY019802 | C(Cc1nccs1)C(O)=O |
| 8 | A0954 | C(C(O)=O)c1csc(N)n1 |
| 9 | STK891980 | C(COc1ccccc1)C([O-])=O |
| 10 | STL193409 | C(CN1C=CC=CC1=O)C([O-])=O |
| 11 | Z1171978885 | C(C([O-])=O)c1cc2ccccc2o1 |
| 12 | STL163377 | C(C([O-])=O)c1nc2ccccc2[nH]1 |
| 13 | Z1333043518 | C(C([O-])=O)n1cc2ccccc2n1 |
| 14 | F1913-0024 | C(C([O-])=O)c1nc2ccccc2o1 |
| 15 | STL124046 | C(C([O-])=O)NC(c1cccnc1)=O |
| 16 | STK526633 | C(CC(c1cccs1)=O)C([O-])=O |
| 17 | STL190846 | C(C([O-])=O)c1cccc2cccnc12 |
| 18 | STK695471 | C(Cc1cn2cccnc2n1)C([O-])=O |
| 19 | EN300-14584 | C(Cn1cnc2ccccc12)C(O)=O |
| 20 | EN300-70527 | C(C(O)=O)N1Cc2ccccc2C1=O |
| 21 | STK504141 | C(C([O-])=O)Nc1ccc2nncn2n1 |
| 22 | STK353430 | C(Cc1nc2ncccn2n1)C(O)=O |
| 23 | Z188924494 | C(Cc1ccc2c(c1)OCO2)C([O-])=O |
| 24 | IF_03_114 | C(CC(Nc1ccccc1)=O)C(O)=O |
| 25 | STK723995 | C(C([O-])=O)Nc1c2ccccc2ncn1 |
| 26 | MS-2504 | C(C([O-])=O)Oc1cnc2ccccc2n1 |
| 27 | STK392449 | C(C([O-])=O)N1C=Nc2ccccc2C1=O |
| 28 | EN300-188561 | C(C(O)=O)N1C=Cc2ccccc2C1=O |
| 29 | STK738252 | C(CN1C(=O)Oc2ccccc12)C([O-])=O |
| 30 | Z57983005 | C(C1C(Nc2ccccc2O1)=O)C(O)=O |
| 31 | STK447336 | Cc1cc(NC(CCC([O-])=O)=O)nn1C |
| 32 | CDS014372 | C1C[C@H](C(N)=O)N(C1)C(CCC([O-])=O)=O |
| 33 | Z126932466 | C(Cc1cnn(c1)c1ccccc1)C(O)=O |
| 34 | STK500480 | C(Cc1nc(c2ccccc2)no1)C([O-])=O |
| 35 | TS-00478 | C(Cc1nnc(c2ccccc2)o1)C([O-])=O |
| 36 | STL310201 | C(CC([O-])=O)C1C(N2C=CC=CC2=NN=1)=O |
| 37 | AL-291/37197008 | C(Cn1nc(c2ccccc2)nn1)C(O)=O |
| 38 | STK099228 | C(CC(N1CCc2ccccc12)=O)C([O-])=O |
| 39 | Z166605066 | C(CN1C(c2ccccc2N=N1)=O)C(O)=O |
| 40 | STK946249 | C(CN1C(c2ccccc2S1)=O)C([O-])=O |
| 41 | EN300-00843 | CC1(C)Cc2cccc(c2O1)OCC(O)=O |
| 42 | EN300-35451 | CC1=C(CCC([O-])=O)C(NC(=N1)SC)=O |
| 43 | Z1259161843 | C(C([O-])=O)Nc1ccc(cc1)S(N)(=O)=O |
| 44 | EN300-10660 | Cc1c2C(N(CC([O-])=O)C=Nc2sc1C)=O |
| 45 | IF_04_062 | C(Cc1nc2cccc3cccc(c23)[nH]1)C(O)=O |
| 46 | IF_03_115 | C(CC(Nc1cccc2ccccc12)=O)C(O)=O |
| 47 | Z90120424 | C(C(O)=O)N1C=Nc2c3ccccc3oc2C1=O |
| 48 | EN300-137714 | C(CC(c1ccc2c(CCC(N2)=O)c1)=O)C(O)=O |
| 49 | Z102925466 | C(CN1C(c2ccccc2S1(=O)=O)=O)C(O)=O |
| 50 | IF_01_187 | C(Cc1ccccc1OCc1ccccn1)C(O)=O |
| 51 | STK925744 | CC1=CC(=O)Oc2cc(c(CCC([O-])=O)cc12)OC |
| 52 | STK682580 | C1CCn2c(C1)c(C#N)c1c2C(N(CC(O)=O)C=N1)=O |
| 53 | KM09821 | C(CC(O)=O)CN1CCC(=CC1)c1ccccc1.[Cl] |
| 54 | STK545027 | CC(C(O)=O)N1C(C(=Cc2ccc(cc2)OC)SC1=O)=O |
| 55 | Z1682077208 | C1CCN(C1)S(c1ccc(cc1)C(CCC(O)=O)=O)(=O)=O |
| 56 | 79148212 | C(Cc1nc2ccccc2n1c1cc(C(N)=O)sc1)C(O)=O |
| 57 | MS-2054 | C(C(Nc1nnc(SCC(O)=O)s1)=O)Oc1ccccc1 |
| 58 | Z609944470 | C(Cc1nc2ccccc2c2nc(Cc3ccccc3)nn12)C([O-])=O |
| 59 | Z1373577255 | C(CC(c1cccc(c1)NS(c1c[nH]c2c1cccn2)(=O)=O)=O)C(O)=O |
| 60 | Z1413959119 | Cc1ccc(cc1S(Nc1cccc(c1)C(CCC(O)=O)=O)(=O)=O)C(O)=O |
| 61 | HGA-4028-0036-50 | C1CN(CCC1c1ccc(cc1)[Cl])C(c1ccc(C(NCC(O)=O)=O)nc1)=O |
| 62 | CAZ-0010-0660-20 | C1CN(CCC1N1C(=O)Oc2ccccc12)S(c1ccc(cc1)C(NCC(O)=O)=O)(=O)=O |
| 63 | Z1373577229 | C1CCN(C1)C(c1cc(c[nH]1)S(Nc1cccc(c1)C(CCC(O)=O)=O)(=O)=O)=O |
| 64 | Z1413958135 | C(CC(c1cccc(c1)NS(c1ccc(c(c1)C(O)=O)[Cl])(=O)=O)=O)C(O)=O |
| 65 | Z1413958985 | C(CC(c1cccc(c1)NS(c1cc(cc(c1)F)C(O)=O)(=O)=O)=O)C(O)=O |
| 66 | Z1413959119 | Cc1ccc(cc1S(Nc1cccc(c1)C(CCC(O)=O)=O)(=O)=O)C(O)=O |
| 67 | Z1413958389 | CS(=O)(=O)NC=1C=CC=C(C1)C(=O)CCC(=O)O |
| 68 | Z1686553089 | OC(=O)CCC(=O)C=1C=CC=C(NS(=O)(=O)C2CC2)C1 |
| 69 | Z1413958377 | OC(=O)CCC(=O)C=1C=CC=C(NS(=O)(=O)N2CCCC2)C1 |
| 70 | Z1715629969 | CC1CCCN1S(=O)(=O)NC=2C=CC=C(C2)C(=O)CCC(=O)O |
| 71 | Z1783325602 | OC(=O)CCC(=O)C=1C=CC=C(NS(=O)(=O)C2=CC=CO2)C1 |
| 72 | Z2067509115 | OC(=O)CCC(=O)C=1C=CC=C(NS(=O)(=O)C=2C=CSC2)C1 |
| 73 | Z1413958278 | OC(=O)CCC(=O)C=1C=CC=C(NS(=O)(=O)C=2C=NNC2)C1 |
| 74 | Z1413957763 | OC(=O)CCC(=O)C=1C=CC=C(NS(=O)(=O)C2=CNC=N2)C1 |
| 75 | Z1413959097 | OC(=O)CCC(=O)C=1C=CC=C(NS(=O)(=O)N2CCCCC2)C1 |
| 76 | Z1373577222 | OC(=O)CCC(=O)C=1C=CC=C(NS(=O)(=O)C=2C=CC=CC2)C1 |
| 77 | Z1373577090 | OC(=O)CCC(=O)C=1C=CC=C(NS(=O)(=O)C=2C=CC=3C=CC=CC3C2)C1 |
| 78 | Z1413957848 | OC(=O)CCC(=O)C=1C=CC=C(NS(=O)(=O)C=2C=CC=C3C=CC=CC23)C1 |
| 79 | Z2010369233 | OC(=O)CCC(=O)C=1C=CC=C(NS(=O)(=O)C=2C=NC=3C=CC=CC3C2)C1 |
| 80 | Z1373577254 | OC(=O)CCC(=O)C=1C=CC=C(NS(=O)(=O)C=2C=CC=3CCCC3C2)C1 |
| 81 | Z1783325595 | OC(=O)CCC(=O)C=1C=CC=C(NS(=O)(=O)C2=CN=C3CCCN23)C1 |
| 82 | Z1738053529 | CN1N=CC=2C=C(C=NC12)S(=O)(=O)NC=3C=CC=C(C3)C(=O)CCC(=O)O |

**Table S2: Matrix Screen results**

| Compound # | KD (µM) | | | | | | | | | | |
| --- | --- | --- | --- | --- | --- | --- | --- | --- | --- | --- | --- |
|  | USP3 | USP5 | USP13 | USP16 | USP20 | USP33 | USP39 | USP49 | USP51 | BRAP | HDAC6 |
| 1 | 121 | 1000 | 1000 | 96.1 | 707 | 115 | 180 | 283 | 1000 | 73.9 | 1000 |
| 2 | 1000 | 1000 | 98.2 | 1000 | 182 | 1000 | 1000 | 56 | 1000 | 74.4 | 1000 |
| 3 | 1000 | 1000 | 1000 | 1000 | 1000 | 1000 | 1000 | 1000 | 1000 | 1000 | 1000 |
| 4 | 178 | 1000 | 1000 | 429 | 1000 | 1000 | 1000 | 172 | 1000 | 82.9 | 1000 |
| 5 | 855 | 1000 | 616 | 1000 | 574 | 1000 | 1000 | 1000 | 1000 | 1000 | 1000 |
| 6 | 1000 | 1000 | 1000 | 1000 | 1000 | 1000 | 1000 | 1000 | 1000 | 1000 | 1000 |
| 7 | 173 | 1000 | 547 | 331 | 439 | 808 | 1000 | 1000 | 1000 | 1000 | 1000 |
| 8 | 1000 | 1000 | 1000 | 1000 | 895 | 1000 | 1000 | 1000 | 1000 | 1000 | 1000 |
| 9 | 1000 | 1000 | 640 | 1000 | 523 | 724 | 1000 | 1000 | 1000 | 1000 | 1000 |
| 10 | 1000 | 1000 | 1000 | 1000 | 1000 | 1000 | 1000 | 1000 | 1000 | 1000 | 1000 |
| 11 | 77.1 | 1000 | 472 | 265 | 472 | 1000 | 1000 | 1000 | 1000 | 1000 | 1000 |
| 12 | 1000 | 1000 | 1000 | 1000 | 1000 | 1000 | 1000 | 1000 | 1000 | 1000 | 1000 |
| 13 | 519 | 1000 | 1000 | 1000 | 715 | 268 | 665 | 1000 | 1000 | 411 | 1000 |
| 14 | 1000 | 1000 | 278 | 1000 | 403 | 368 | 1000 | 1000 | 1000 | 1000 | 1000 |
| 15 | 63.2 | 1000 | 1000 | 198 | 1000 | 514 | 1000 | 1000 | 1000 | 1000 | 1000 |
| 16 | 195 | 1000 | 194 | 1000 | 272 | 1000 | 1000 | 1000 | 1000 | 1000 | 555.4 |
| 17 | 1000 | 1000 | 1000 | 1000 | 1000 | 1000 | 1000 | 1000 | 1000 | 1000 | 1000 |
| 18 | 1000 | 1000 | 833 | 1000 | 956 | 1000 | 1000 | 1000 | 1000 | 416 | 1000 |
| 19 | 1000 | 1000 | 1000 | 1000 | 1000 | 1000 | 1000 | 1000 | 1000 | 1000 | 1000 |
| 20 | 758 | 1000 | 1000 | 1000 | 1000 | 1000 | 1000 | 525 | 1000 | 685 | 1000 |
| 21 | 1000 | 1000 | 780 | 1000 | 882 | 1000 | 1000 | 1000 | 1000 | 1000 | 869 |
| 22 | 1000 | 1000 | 118 | 1000 | 110 | 1000 | 1000 | 1000 | 1000 | 1000 | 1000 |
| 23 | 1000 | 1000 | 1000 | 1000 | 1000 | 1000 | 1000 | 1000 | 1000 | 1000 | 1000 |
| 24 | 1000 | 1000 | 872 | 1000 | 817 | 1000 | 1000 | 1000 | 1000 | 1000 | 1000 |
| 25 | 191 | 1000 | 1000 | 603 | 1000 | 1000 | 1000 | 1000 | 1000 | 1000 | 1000 |
| 26 | 1000 | 1000 | 1000 | 1000 | 1000 | 1000 | 1000 | 1000 | 1000 | 885 | 1000 |
| 27 | 328 | 893 | 860 | 456 | 835 | 1000 | 1000 | 1000 | 1000 | 1000 | 1000 |
| 28 | 249 | 1000 | 228 | 1000 | 224 | 1000 | 1000 | 1000 | 1000 | 1000 | 698.4 |
| 29 | 817 | 1000 | 1000 | 981 | 1000 | 1000 | 1000 | 1000 | 1000 | 993 | 1000 |
| 30 | 1000 | 1000 | 306 | 1000 | 271 | 1000 | 1000 | 1000 | 1000 | 1000 | 1000 |
| 31 | 201 | 1000 | 660 | 193 | 777 | 1000 | 1000 | 1000 | 1000 | 1000 | 1000 |
| 32 | 165 | 265 | 1000 | 154 | 1000 | 1000 | 1000 | 399 | 1000 | 462 | 565 |
| 33 | 1000 | 1000 | 1000 | 1000 | 1000 | 1000 | 1000 | 661 | 1000 | 1000 | 1000 |
| 34 | 1000 | 1000 | 128 | 1000 | 119 | 1000 | 1000 | 1000 | 1000 | 1000 | 1000 |
| 35 | 1000 | 831 | 220 | 1000 | 64.7 | 1000 | 1000 | 1000 | 1000 | 340 | 1000 |
| 36 | 1000 | 1000 | 1000 | 1000 | 1000 | 1000 | 1000 | 1000 | 1000 | 1000 | 1000 |
| 37 | 146 | 1000 | 950 | 482 | 677 | 1000 | 1000 | 1000 | 1000 | 1000 | 231.8 |
| 38 | 1000 | 1000 | 294 | 1000 | 291 | 1000 | 1000 | 1000 | 1000 | 646 | 95.6 |
| 39 | 102 | 1000 | 1000 | 1000 | 1000 | 851 | 1000 | 1000 | 1000 | 1000 | 331.4 |
| 40 | 1000 | 1000 | 1000 | 1000 | 1000 | 1000 | 1000 | 1000 | 1000 | 1000 | 1000 |
| 41 | 583 | 1000 | 1000 | 671 | 1000 | 1000 | 1000 | 1000 | 1000 | 1000 | 1000 |
| 42 | 51.8 | 1000 | 1000 | 118 | 814 | 211 | 389 | 1000 | 1000 | 1000 | 1000 |
| 43 | 620 | 1000 | 1000 | 428 | 1000 | 1000 | 1000 | 1000 | 1000 | 1000 | 1000 |
| 44 | 290 | 238 | 1000 | 579 | 674 | 1000 | 1000 | 1000 | 1000 | 315 | 187.6 |
| 45 | 1000 | 1000 | 1000 | 1000 | 1000 | 1000 | 1000 | 1000 | 1000 | 1000 | 1000 |
| 46 | 1000 | 1000 | 1000 | 1000 | 1000 | 1000 | 1000 | 1000 | 1000 | 66 | 198.8 |
| 47 | 1000 | 312 | 1000 | 344 | 426 | 1000 | 1000 | 1000 | 1000 | 253 | 247 |
| 48 | 140 | 532 | 1000 | 167 | 821 | 1000 | 1000 | 99.9 | 1000 | 126 | 103.6 |
| 49 | 1000 | 1000 | 1000 | 1000 | 1000 | 1000 | 1000 | 1000 | 1000 | 1000 | 1000 |
| 50 | 49.5 | 1000 | 365 | 48.2 | 249 | 1000 | 1000 | 1000 | 1000 | 483 | 1000 |
| 51 | 354 | 1000 | 867 | 535 | 465 | 1000 | 1000 | 86.2 | 1000 | 71.6 | 343.2 |
| 52 | 603 | 207 | 1000 | 403 | 552 | 1000 | 1000 | 1000 | 1000 | 215 | 105.4 |
| 53 | 1000 | 561 | 594 | 1000 | 380 | 1000 | 1000 | 1000 | 1000 | 1000 | 1000 |
| 54 | 441 | 1000 | 1000 | 116 | 457 | 1000 | 1000 | 1000 | 1000 | 1000 | 1000 |
| 55 | 91.5 | 117 | 1000 | 139 | 798 | 1000 | 1000 | 1000 | 1000 | 340 | 51.54 |
| 56 | 1000 | 1000 | 754 | 1000 | 464 | 1000 | 1000 | 1000 | 1000 | 1000 | 1000 |
| 57 | 219 | 315 | 745 | 372 | 401 | 1000 | 1000 | 1000 | 1000 | 703 | 273.4 |
| 58 | 238 | 847 | 1000 | 285 | 1000 | 1000 | 1000 | 1000 | 1000 | 1000 | 187.4 |
| 59 | 19.1 | 168 | 1000 | 69.8 | 706 | 1000 | 1000 | 1000 | 1000 | 413 | 211.6 |
| 60 | 283 | 1000 | 1000 | 114 | 1000 | 537 | 942 | 187 | 1000 | 234 | 1000 |
| 61 | 344 | 71.1 | 642 | 436 | 430 | 1000 | 1000 | 1000 | 1000 | 537 | 415 |
| 62 | 216 | 127 | 1000 | 536 | 1000 | 1000 | 1000 | 1000 | 1000 | 1000 | 234.1 |
| Ub | 4.83 | 3.01 | 1000 | 8.55 | 1000 | 1000 | 1000 | 1000 | 1000 | 35 | 9.134 |
| Ub-RGGdel | 1000 | 1000 | 1000 | 1000 | 1000 | 1000 | 1000 | 1000 | 1000 | 1000 | 1000 |

**Table S3: PLGS filtering parameters**

| **Parameter** | Min. intensity | Min. sequence length | Max. sequence length | Min. products per amino acid | Minimum score | Max. MH+ Error (ppm) |
| --- | --- | --- | --- | --- | --- | --- |
| **Value** | 20000 | 4 | 26 | 0.12 | 7 | 5 |

**Figure S1: HDX sequence coverage of USP3^1-131^**

A total of 59 peptides yielded 93.9% sequence coverage and a redundancy of 5.24. Black rectangles aligned to sequence represent peptides. Several peptides including 52-86 (length 35) and 53-86 (length 34) were manually added after PLGS filtering to obtain coverage from 53-60.

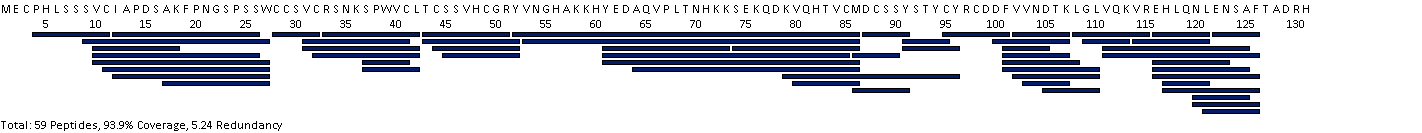

**Figure S2: Differential HDX heatmaps**

Differential fractional uptake is displayed as a per-residue averaged heatmap aligned to USP3 primary sequence for USP3 Zf-UBD in complex with **59** at 1:10 (top) and 1:20 (bottom) . The highest magnitude D decreases (-5 %) are blue and D increases (+5%) are red. A limitation of this heatmap illustration (DynamX 3.0) is that is does not factor in the associated error for each residue. Please refer to Figure 4 for a peptide-level differential HDX bar-plot with associated error.

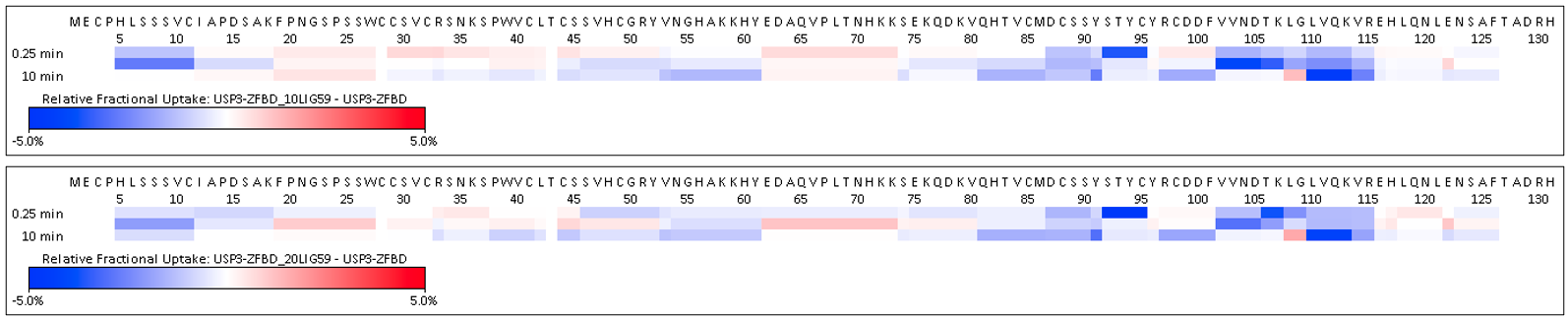

**Figure S3: Ligand RMSD over 100 ns MD simulation**

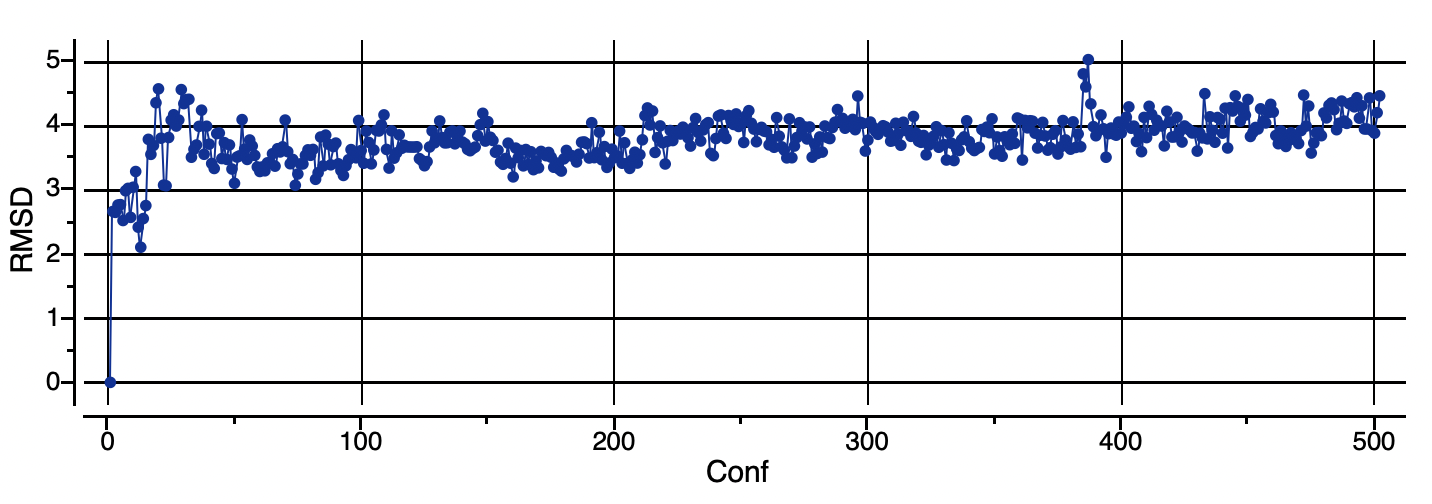
